# Organizing laboratory information to analyze the reproducibility of experimental workflows

**DOI:** 10.1101/2022.04.05.487214

**Authors:** Jean Peccoud, Derek Johnson, Samuel Peccoud, Julia Setchell, Wen Zhou

## Abstract

Reproducibility is the cornerstone of scientific experiments. Assessing the reproducibility of an experiment requires analyzing the contribution of different factors to the variation of the observed data. Suitable data structures need to be defined prior to the data collection effort so that data associated with these factors can be recorded and associated with observations of the variable of interest. The resulting datasets can be analyzed statistically to estimate the effect of experimental factors on the observed data using ANOVA models. Custom data structures to document the execution of experimental workflows are defined in a research data management system. The data produced by multiple repetitions of a plasmid purification process and a cell culture process are analyzed using the Kruskal–Wallis H-test to identify factors contributing to their variation. Repetitions of the plasmid purification process do not lead to significant differences in extraction yields. Statistically significant differences in plasmid solution purity are identified but the differences are small enough that are not biologically relevant. The maintenance of two cell lines over many generations leads to similar datasets. However, different media preparations appear to influence the variation of cell viability and harvested cell counts in unexpected ways that may be the indirect expression of hidden effects not captured in the data structure.

## Introduction

Reproducibility is the difference between an anecdotal observation and the result of a research project [1–3]. Scientific results are expected to be reproducible. The word “reproducibility” can have different meanings in different contexts. It may refer to the reproducibility of computational processes whereas “repeatability” refers to the possibility of repeating a data collection process while reaching similar conclusions [4]. Since the use of these two terms has been a source of confusion for years [5], we use reproducibility to refer to the combination of experimental and computational steps that makes it possible to repeat a data collection and analysis process so that different repetitions lead to identical conclusions.

Yet, the scientific community has come to recognize that it is struggling with serious reproducibility issues. Most scientists have failed to reproduce somebody’s else experiments and even their own results [6]. As a well-known example, using the invaluable Reproducibility Project: Psychology (RP:P) [7], the Open Science Collaboration [8] reported that 64% of the replication studies fail to detect statistically significant results aligning with the original research and 53% of 95%-confidence intervals in the replication studies do not cover the claimed signals from original studies. The problem is so prevalent that the community is wondering if the scientific enterprise is facing a reproducibility crisis [9,10] that may undermine its credibility [11,12].

The reproducibility of life science research has also been documented in detail. For example, the improper authentication of cell lines is a widespread problem that compromises the reproducibility of results described in millions of peer-reviewed publications [13–17]. It is estimated that more than half of the preclinical research budgets are spent on projects that failed to produce reproducible results [18]. The cost of poor reproducibility is also felt at the level of individual labs. While it may not be perceived as a direct waste of scarce financial resources, poor reproducibility translates into delayed publications, inability to maintain expertise within a group, lower number of citations, and, in the most dramatic cases, scientific integrity investigations. Even though it may be difficult to estimate the economic impact of these consequences of poor reproducibility at the individual level, there is little doubt that they translate into opportunity costs in terms of the lower scientific impact of ongoing research projects and increased difficulty getting access to funding to support future research projects [19, 20]. Therefore, the motivation for improving reproducibility should not be limited by a need to comply with requirements imposed by funding agencies or journal editors. It can be justified by the need to optimize the use of existing financial resources.

Awareness of the individual benefits of greater reproducibility is necessary to design research plans and experiments with reproducibility in mind instead of considering reproducibility as an afterthought dictated by compliance requirements. This change of perspective is necessary for analyzing the reproducibility of an experiment and calls for an analysis of the factors that might contribute to the variation of the observed data. Controlled factors in an experiment are expected to cause the effect observed in data. For example, different plasmid designs may lead to different levels of gene expression. However, extraneous factors may influence the observed data in unexpected ways and lead to spurious discoveries. For example, different operators performing the experiments, different versions of a protocol, or different kits used to perform gene expression assays may have unanticipated effects on the outcomes of an experiment. In order to disentangle how do different factors contribute to the variation of data, it is necessary to plan and document the data collection discretely, which is conducive to associating the controlled variables of interest with the observed variation in experimental outcomes correctly. Without this additional data, spurious discoveries might cause misleading scientific conclusions. For example, it is likely the controlled factor in the experiment (i.e. plasmid design) fails to completely explain the variations in data (i.e. gene expression data). The process of fully documenting an experiment by retaining observations of the factor of interest as well as side information describing experimental parameters can be greatly facilitated by properly configuring a data management system ahead of the experiment.

Laboratory information management systems (LIMS) are software tools used by laboratories to manage data related to laboratory operations [21–23]. They can be used to manage inventories, Standard Operating Procedures, data produced by instruments, and more advanced workflows. While most LIMS applications have been designed as enterprise software products designed for corporate laboratories, the synthetic biology community has developed software tools supporting its data-driven research workflows [24, 25].

One of the limitations of current LIMS systems is their focus on laboratory operations rather than data analytics. Their ability to manage complex datasets is limited which makes them ill-suited to support advanced data analytics pipelines. Other software applications have been designed to manage experimental datasets that could be used for the analysis of data [26]. In order to avoid a possible disconnect between the requirements of data analysis of workflows and the data collected during laboratory operations, GenoFAB provides an environment allowing laboratory operators to plan experimental workflows by developing structured data types suitable to support data analytics processes [27].

Here, the GenoFAB data model is applied to analyze the reproducibility of two simple laboratory processes. Using a non-parametric statistical test, we analyze the contributions of different factors influencing the variation of data characterizing the performance of a plasmid purification process and a cell culture process.

## Methods

### Data structure

Data were managed in the GenoFAB LIMS. GenoFAB organizes data in a customizable database of objects using three layers of abstraction.

*Catalog types* are defined by specifying data fields used to describe this class of records. Catalog type definitions include a user-defined key used to generate human-readable identifiers by serializing all the items in this catalog category. Catalog Type definitions also include user-defined data fields selected among supported field types such as short text, rich text, durations, numbers with units, currencies, URL, file attachment, and more. Data fields are organized in two columns corresponding to the data associated with catalog entries and data associated with catalog items. The entry fields are parameters that describe how items should be obtained. A product reference number would be an entry-level field for an instrument whereas a serial number or a warranty expiration date would be item-level fields for an instrument. *Catalog entries* are data objects within a catalog type. Catalog entries are defined by assigning values to the entry-level data fields.

*Catalog items* are instances of catalog entries corresponding to a realization of a catalog entry. They are defined by setting the value of the item-level data fields.

For example, “MCUL: HEK293T/17” is an entry in the “Cell Culture” catalog type that is used to manage the data of all the cultures of this cell line. It is defined by specifying the cell line, the growth media, and other parameters. #MCUL3271 refers to a particular culture that is defined by specifying data such as start date, end date, cell viability, or the number of cells harvested. In addition to this vertical organization from generic classes of objects to instances of specific objects, the GenoFAB database makes it possible to link catalog entries to another. The links are directional. They indicate how records are derived from each other. For example, a cell culture type will include a link to a media and to another culture used as inoculum to specify the media and culture used to start a new culture. Links between catalog records ensure the consistency of the links at the entry and item level. For example, the catalog entry for “DMEM 10%FCS”, a growth media, would include two links to “DMEM” and “FCS”, the two supplies used to make the media. Declaring these links at the catalog entry-level creates links at the item level allowing users to specify which DMEM and FCS orders were used to prepare a batch of “DMEM 10% FCS”.

### Plasmid minipreps

Plasmids were transfected in One Shot^®^ Stbl3™ Chemically Competent *E. coli* (Thermofisher). An individual colony was grown in Luria Bertani (LB) growth medium in the presence of ampicillin. The culture was frozen by mixing 500 ul of culture and 500 ul of 50% (v/v) glycerol/water solution. The glycerol stocks were kept at −80C.

Bacterial cultures were started by inoculating 5 ml LB in the presence of ampicillin (100 ug/ml). After 16 to 18 hours, plasmids were extracted from 1 ml of culture using the ZymoPURE Plasmid Miniprep Kit (Zymo Research) according to the manufacturer’s instructions. The plasmids were recovered in 50 ul of elution buffer.

The concentration and quality of the plasmid solutions were determined by diluting 1 ul of plasmid solution in 4 ul of Tris EDTA buffer and loading 3 ul of the diluted solution on a μCuvette G1 (Eppendorf). Optical densities were measured at 260 nm and 280 nm using a BioSpectrophotometer and Fluorimeter (Eppendorf). Concentrations were derived from optical densities by the instrument software.

### Cell cultures

The adherent cell line HEK293T/17 (ATCC) was grown in 15 ml of DMEM 10% FCS media composed of Dulbecco’s Modified Eagle Medium (Corning) and 10% Fetal Bovine Serum (Fisher Scientific). The CMV-mEGFP-Puro cell line was produced by transfecting HEK293T/17 with a plasmid expressing the mEGFP gene under the control of the CMV promoter. Stable transformants were selected by growing the cell line in selective media (DMEM 10% FCS 1 μg/ml puromycin). T75 flasks (Corning) were inoculated with 1 to 4 million cells. Cells were harvested when reaching 70-80% convergence. The cells were harvested by trypsinization. The viability and number of cells in the resulting suspension were determined by counting the cells after staining with Trypan Blue.

### Data analysis

Data were analyzed using the SciPy Python library. Plasmid data are available as Supplementary Data S1. Cell culture data are available as Supplementary Data S2

The analysis script is provided as a Jupyter notebook [28] as Supplementary Data S3.

## Results

### Plasmid preparation

We performed minipreps from 5 high-copy number mammalian expression vectors derived from pUC19. Their size ranged from 2.6kb to 7.7kb (Table 1).

**Table 1:** List of plasmids used in this study.

| Plasmid ID | Size (bp) |
| --- | --- |
| DPLA3111 | 2,686 |
| DPLA483 | 4,966 |
| DPLA2104 | 5,223 |
| DPLA2105 | 7,691 |
| DPLA2101 | 6,824 |

We repeated the culture and DNA extraction process three times. Each plasmid solution was analyzed by measuring the optical density (OD) at 260nm and 280nm to estimate the concentration and quality of the plasmid solution. The OD measurement was performed three times for each solution. The spectrometer software converted the optical densities into concentrations in ng/ul using the sample dilution. The instrument also produced the ratio of A260/A280 which is often used as an indication of preparation quality. A low value of this ratio (<1.8) is indicative of the residual presence of reagents used in the DNA extraction process. High A260/A280 ratios are not indicative of any specific issue.

The Supplementary Data S1 provides all the raw data organized by replicate. This file also shows the standard deviation and coefficient of variation of the A260 measurements. For most samples, the variation of the OD measurement is less than 10%. Two samples (#SPLA3ll6, #SPLA3117) had a coefficient of variation of A260 observations around 20%. The A260/A280 data have CV less than 5% with some samples having CV less than 1%.

All the subsequent analysis focuses on the variation across mean of A260 and A260/A280 values collected on individual plasmid preparations.

A rapid examination of Supplementary Data S1 indicates that one plasmid seems to lead to higher concentrations (DPLA2105 while plasmid DPLA3111 leads to lower plasmid concentrations. Boxplots of the plasmid concentrations (Fig. 1A) and A260/A280 distributions (Fig. 1B) display the data’s empirical distributions. These figures suggest that the distributions of plasmid concentration vary across the plasmid types. However, the A260/A280 distributions do not seem to be influenced by the plasmid as much as the concentration.

**Figure 1:**
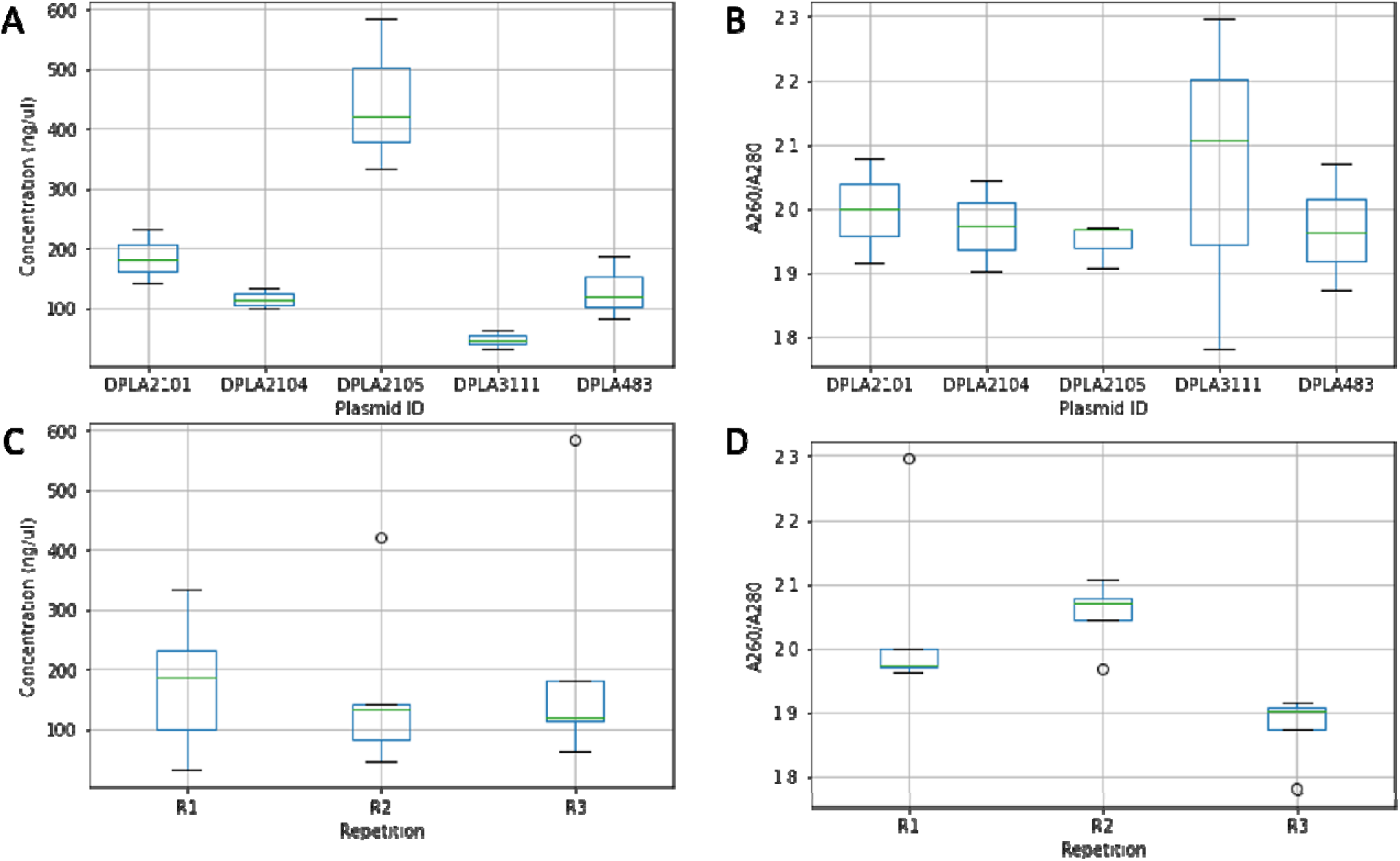
Reproducibility of the plasmid purification process. Boxplots showing the variation of plasmid concentration (A, C) and A260/A280 ratio (B, D). The process variability is analyzed across two model parameters: the plasmids (A, B) and the repetition (C, D).

Next, we generated boxplots to examine the variation across repetitions. Instead of looking at three replicates of the plasmid purification process for each plasmid, we compared concentrations and A260/A280 values for all the five plasmids for each of the replicates. Even though all the replicates are supposed to be identical, it is possible that some purification processes went better than others leading to differences across all the plasmids. While the concentration values seem homogeneous across the three replicates (Fig. 1C), there are noticeable differences in A260/A280 values (Fig. 1D).

The boxplots help articulate a statistical hypothesis on the systematic deviation across plasmids. This is essentially a one-way analysis of variance (ANOVA). For such a hypothesis, many tests are available, including the rank-based Kruskal-Wallis H-test that is robust against outliers and the lack of normality in the data [29]. The test compares different groups of samples and tests the null hypothesis that they originate from the same distribution. A small p-value indicates statistically significant differences across different groups.

*We* performed four H-tests to investigate the plasmid and repetition effects on the concentration and A260/A280 values (Table 2). At the nominal significance level of 0.05, with the Bonferroni correction of the multiplicity [30], the tests confirm that there is a significant repetition effect on the A260/A280 values while the plasmid effect on the concentration values is substantial but less significant at the 0.05 level. In other words, the nature of the plasmid affects the yield of the purification process somehow but not the quality of the plasmid solutions. The three repetitions lead to similar plasmid concentrations but to different A260/A280 values. However, since the values are all greater than 1.8, the statistically significant difference observed is probably not biologically meaningful. This suggests that the plasmid purification process is not necessarily irreproducible since the variation across repetitions is unlikely to be biologically relevant.

**Table 2:** p-values ofH tests for the plasmid minipreps dataset. Adjusted for the multiplicity of hypotheses in the table, the plasmid effect on concentration is significant at the nominal significance level of 0.1 but not at the level of 0.05, the repetition effect on A260/A280 ratio is significant at both the nominal levels of 0.1 and 0.05.

| Variable/Effect | Plasmid | Repetition |
| --- | --- | --- |
| Concentration | 0.016* | 0.932 |
| A260/A280 | 0.814 | 0.007** |

### Cell culture

We maintained two mammalian cell lines derived from HEK293. The parent line is HEK293T/17. The second line stably expresses a reporter gene and is maintained in selective media.

Over the course of several months, we recorded the number of cells used to seed the culture, the duration of the culture in numbers of days between seeding and harvesting, the viability of the culture at harvest time, and the number of cells harvested. In addition, we recorded the media preparation used for each culture.

A subset of these data is available as supplementary data S2. The data set has been cleaned to remove incomplete entries. The cleaned dataset includes 32 entries of the original cell line and 37 entries for the transformed cell line.

Considering that the transformed cell line is maintained in selective media and expresses two transgenes at a high level, it is possible that its behavior would differ significantly from the properties of the original cell lines. Differences in fitness could translate into lower viability rates, smaller numbers of cells at harvest time, or different times to reach convergence.

We compared the two cell lines visually (Fig. 2 A-C) and also conducted the H-test on the effect of cell lines (Table 3). These comparisons demonstrate that the behavior of the two cell lines is remarkably similar. There is no difference in the viability, cell count, or culture duration that can be identified between the two groups of cultures.

**Figure 2:**
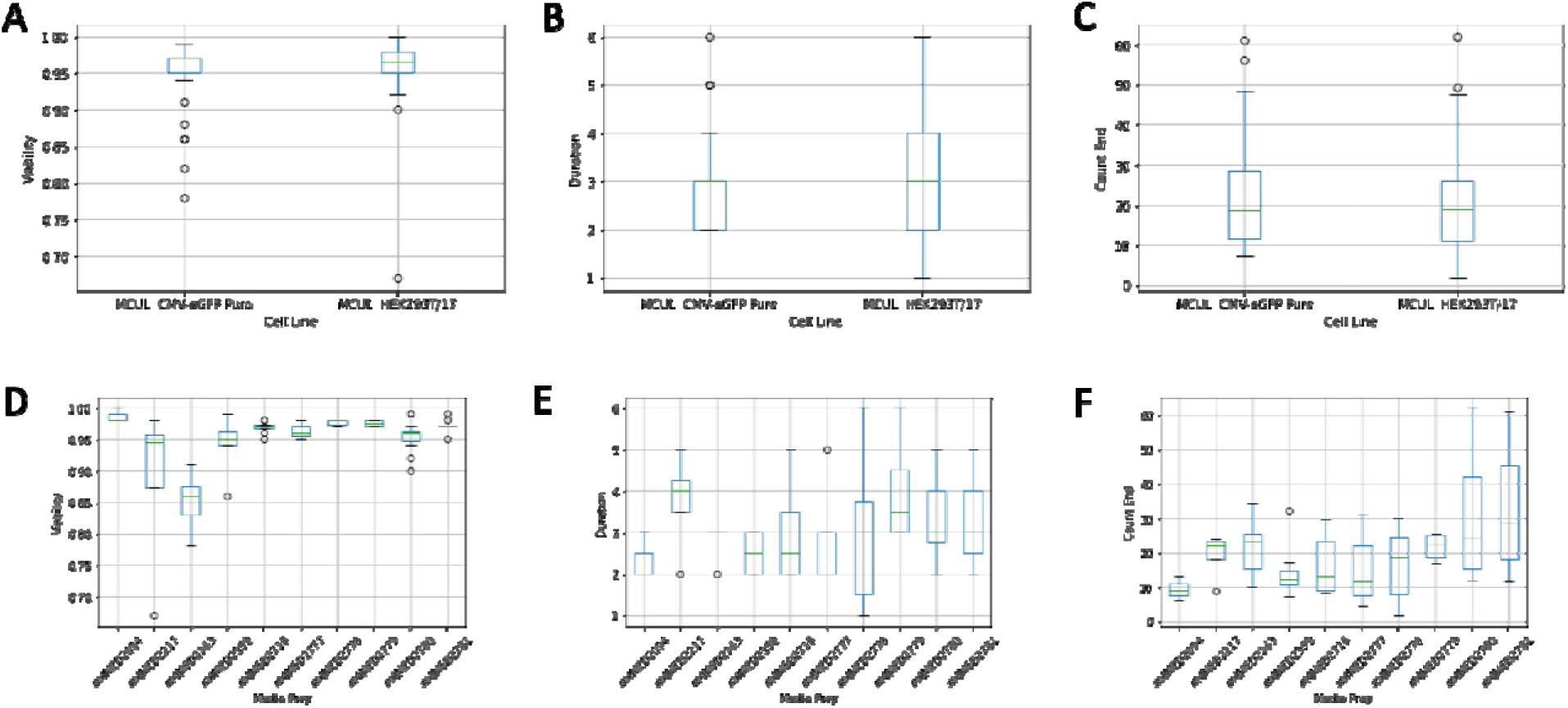
Reproducibility of cell cultures. Boxplots showing the variation of the viability (A, D), number of days until achieving 80% convergence (B, E), and number of cells at harvest time (C, F) in a set of 70 cultures of adherent mammalian cells. The process variability is analyzed across two model parameters: the cell line (A-C) and the media preparation (D-F).

**Table 3:** p-values ofH tests for cell culture dataset. Adjusted for the multiplicity of hypotheses in the table, the media preparation influences on viability significantly at both nominal significance levels of 0.1 and 0.05.

| Variable/Effect | Cell Line | Media Prep. |
| --- | --- | --- |
| Viability | 0.458 | 0.0001*** |
| Duration | 0.984 | 0.332 |
| Cell Count End | 0.523 | 0.031* |

Then we explored the possibility that the media preparation could have an impact on cell growth. Media are prepared in batches of 550 ml by combining 500 ml of base media with 50 ml of fetal calf serum. A media preparation lasts about a month or so. Differences between media preparations could include different lots of fetal calf-serum or simply differences of “freshness” as some preparations may be used faster than others based on the lab workload. Differences between media preparations were analyzed visually (Fig. 2 D-F) and statistically (Table 3). Surprisingly, with the multiplicity of hypotheses corrected at the nominal level of 0.05, a significant difference is observed across media preparation for the viability of the cell culture. Media preparation does not appear to influence the number of days between seed and harvest and the number of cells harvested significantly.

Further analysis would be necessary to interpret this observation. Statistical tests might be sensitive to the existence of indirect effects, statistical dependence among data, and distributional shift [31, 32].

Media preparations correspond to periods of time of approximately 30 days. This may lead the effect associated with the media preparation to be indirect. The media preparation effect may be the indication of a temporal effect rather than an indication of the variability of the media itself. For example, the cell counting process used to calculate the culture viability may have been performed differently during periods of time corresponding to specific media preparation.

## Discussion

### Analysis of variance

Reproducibility can be a misleading concept. A reproducible experiment is not an experiment that leads to identical data every time due to randomness. They always present some level of variation. As a result, assessing the reproducibility of an experiment requires understanding the factors that contribute to the variation of the observed variables. This can be interpreted as an ANOVA problem.

The process of analyzing the variance components of a dataset starts with building an appropriate statistical model that includes factors potentially responsible for the data variation [33]. Then statistical inference is drawn to quantify the contributions of these factors to the total variation.

Experimentalists need to understand that assessing the reproducibility of a dataset needs to be planned when the experiment is designed. In particular, they need to consider what factors might contribute to the variation of the data and record all the information necessary for analysis. The data structure proposed here facilitates this process by identifying different catalog entries associated with different parameter values. The relationships coded in input fields provide another mechanism to identify how factors contribute to data variations. ANOVA models are commonly used in genetics to estimate genetic effects [34]. They typically involve linear combinations of effects such as alleles at different loci, environmental factors as well as interactions effects. The interpretation of ANOVA relies on a few standard statistical assumptions including the normality and homoscedasticity in errors.

Data produced by laboratory workflows like cell cultures or plasmid extraction processes lead to small sample sizes because of the high cost of replicates. It is not always reliable to assume normality in data given the small sample size. The rank-based Kruskal-Wallis H-test offers a robust and simple way to facilitate the one-way analysis of variance for one factor at a time [29]. The test does not require a balanced design. As a one-way ANOVA test, the H-test only reflects the existence of differences among different groups, but it does not indicate what groups stand out. It is also important to keep in mind that small sample sizes can lead to falsepositive results.

The plasmid data set would probably benefit from additional repetitions to investigate the differences observed between plasmids. It may also be worth trying to identify if the differences observed can be inputted to a single outlying plasmid by performing the H-test using a jackknife resampling strategy whereby subsets of the data are analyzed after elimination of the data associated with one plasmid at a time.

#### Fixed and Random Effects

Our analysis of these two datasets included two effects each [35–37]. The analysis of the plasmid purification data considered the plasmid and the repetition effects. The analysis of the cell culture data considered the cell line and media preparation effects.

The plasmid and cell line effects are fixed because they can be controlled experimentally. The repetition and the media preparation lead to random effects. We assume that they may contribute to the data variation, but these effects are random rather than control parameters. It is reasonable to assume that different repetitions of the purification process may lead to different results, but we simply do not know what aspect of the purification process might influence the resulting data. It can be the operator, the kit used, the time of the day, the duration of the cell culture, or anything else that might influence the purification process. Similarly, the media preparation might influence the cell culture process in ways that we cannot control.

The reproducibility of an experiment does not mean that the data are identical but that a hypothesis can be consistently rejected with a certain confidence level and the true signal can be estimated in the same directions. The observation that some plasmids consistently lead to higher purification yield is a reproducible observation that can support a rational effort to improve plasmid performance. Differences associated with other fixed effects may indicate a problem that needs to be addressed. For example, observing differences between operators or between instruments used to execute a process is probably indicative of a problem that should be addressed by recalibrating the instrument or training an operator.

Observing statistically significant differences associated with random effects is more problematic. Large random effects might dominate fixed effects and prevent the detection of meaningful fixed effects. Though it might be handled with more sophisticated statistical models, the existence of large random effects always indicates potential subgroup structures, distributional shifts, or even endogeneity. In general, the random effects are challenging to interpret. They may result from hidden fixed effects that should be identified and controlled.

### Increasing reproducibility through virtualization of research processes

Historically, many life science research projects have relied on “semi-quantitative” approaches that compared data on small groups of replicates without proper statistical analysis [38–41]. Reviewers and editors understood that the time and cost of replicating some experiments made it practically challenging to replicate them. In this context, the consistency between the data and a biological model was often considered sufficient to publish results without proper statistical analysis of the result reproducibility.

As the scientific community recognizes the limitations of this traditional approach, it has come to imagine new ways to accelerate research processes and reduce the cost of data points [42]. Outsourcing certain steps to specialized contract research organizations has become very common. Oligonucleotide synthesis, sequencing, and gene synthesis are some of the operations that have been industrialized resulting in faster and cheaper service than it is possible to achieve when performing these operations for individual projects. More recently, many leading academic institutions have invested in the development of biofoundries that are high-throughput facilities capable of executing complex research workflows rather than individual steps [43–48]. This transformation of the research enterprise should positively impact the reproducibility of research results. Automation reduces the number of random effects by standardizing operations in a way that is not possible with manual operators. High throughputs reduce the cost of individual data points and therefore increase statistical power.

While the research enterprise is undergoing its own industrial revolution, the main obstacle to improving the reproducibility of research results may be human factors. Generations of scientists have been trained to execute experiments in their own laboratories. Today, an increasing amount of data is produced by workers at contract research and manufacturing organizations [49, 50] who execute industrial processes. Statistically, a substantial number of efforts have been documented to enhance and assess the reproducibility of high throughput experiments, especially those in genomics and genetics [51–55]. In this context, scientific expertise is increasingly shifting away from the execution of experiments toward the ability to design and analyze complex datasets collected by a global network of service providers.

## Supporting information

S3-Jupyter notebook

## Material Availability

The plasmids and cell cultures are available upon request.

## Data Availability

The raw data and data analysis scripts are available in the article Supplementary Data.

