## Supplementary material for "Organizing laboratory information to analyze the reproducibility of experimental workflows": S3-Jupyter notebook: S3-reproducibility.ipynb

```
# %% [code]
import pandas
import scipy
from scipy import stats
import numpy as np
import matplotlib.pyplot as plt
df = pandas.read_csv('minipreps.csv')
print(df)

# %% [code]
bp1 = df.boxplot(column=['Mean_C'], by='Plasmid ID') 
bp1.set_xlabel("Plasmid ID")
bp1.set_ylabel("Concentration (ng/ul)")
plt.title( '' )
plt.suptitle('')
plt.show()

# %% [code]
bp2 = df.boxplot(column=['Mean_Q'], by='Plasmid ID')
bp2.set_xlabel("Plasmid ID")
bp2.set_ylabel("A260/A280")
plt.title( '' )
plt.suptitle('')
plt.show()

# %% [code]
bp3 = df.boxplot(column=['Mean_C'], by='Repetition') 
bp3.set_xlabel("Repetition")
bp3.set_ylabel("Concentration (ng/ul)")
plt.title( '' )
plt.suptitle('')
plt.show()

# %% [code]
bp4 = df.boxplot(column=['Mean_Q'], by='Repetition')
bp4.set_xlabel("Repetition")
bp4.set_ylabel("A260/A280")
plt.title( '' )
plt.suptitle('')
plt.show()

# %% [code]
scipy.stats.kruskal(*[group["Mean_C"].values for name, group in df.groupby("Repetition")])

# %% [code]
scipy.stats.kruskal(*[group["Mean_C"].values for name, group in df.groupby("Plasmid ID")])

# %% [code]
scipy.stats.kruskal(*[group["Mean_Q"].values for name, group in df.groupby("Repetition")])

# %% [code]
scipy.stats.kruskal(*[group["Mean_Q"].values for name, group in df.groupby("Plasmid ID")])

# %% [code]
df = pandas.read_csv('cells_v4.csv')
print(df)

# %% [code]
bp1 = df.boxplot(column=['Viability'], by='name') 
bp1.set_xlabel("Cell Line")
bp1.set_ylabel("Viability")
plt.title( '' )
plt.suptitle('')
plt.show()
scipy.stats.kruskal(*[group["Viability"].values for name, group in df.groupby("name")])

# %% [code]
bp1 = df.boxplot(column=['Count End'], by='name') 
bp1.set_xlabel("Cell Line")
bp1.set_ylabel("Cell Count End")
plt.title( '' )
plt.suptitle('')
plt.show()
scipy.stats.kruskal(*[group["Count End"].values for name, group in df.groupby("name")])

# %% [code]
bp1 = df.boxplot(column=['Duration'], by='name') 
bp1.set_xlabel("Cell Line")
bp1.set_ylabel("Duration")
plt.title( '' )
plt.suptitle('')
plt.show()
scipy.stats.kruskal(*[group["Duration"].values for name, group in df.groupby("name")])

# %% [code]
bp1 = df.boxplot(column=['Viability'], by='Media') 
bp1.set_xlabel("Media Prep")
bp1.set_ylabel("Viability")
plt.title( '' )
plt.suptitle('')
plt.xticks(rotation = 45)
plt.show()
scipy.stats.kruskal(*[group["Viability"].values for name, group in df.groupby("Media")])

# %% [code]
bp1 = df.boxplot(column=['Count End'], by='Media') 
bp1.set_xlabel("Media Prep")
bp1.set_ylabel("Count End")
plt.xticks(rotation = 45)
plt.title( '' )
plt.suptitle('')
plt.show()
scipy.stats.kruskal(*[group["Count End"].values for name, group in df.groupby("Media")])

# %% [code]
bp1 = df.boxplot(column=['Duration'], by='Media') 
bp1.set_xlabel("Duration")
bp1.set_ylabel("Count End")
plt.xticks(rotation = 45)
plt.title( '' )
plt.suptitle('')
plt.show()
scipy.stats.kruskal(*[group["Duration"].values for name, group in df.groupby("Media")])

# %% [code]
```
